# Proteins containing 6-crossing knot types and their folding pathways

**DOI:** 10.1101/2023.06.16.545156

**Authors:** Maciej Sikora, Erica Flapan, Helen Wong, Pawel Rubach, Wojciech Garstka, Szymon Niewieczerzal, Eric J Rawdon, Joanna I. Sulkowska

## Abstract

Studying complex protein knots can provide new insights into potential knot folding mechanisms and other fundamental aspects of why and how proteins knot. This paper presents results of a systematic analysis of the 3D structure of proteins with 6-crossings knots predicted by the artificial intelligence program AlphaFold 2. Furthermore, using a coarse-grained native based model, we found that three representative proteins can self tie to a 6_3_ knot, the most complex knot found in a protein thus far. Because it is not a twist knot, the 6_3_ knot cannot be folded via a simple mechanism involving the threading of a single loop. Based on successful trajectories for each protein, we determined that the 6_3_ knot is formed after folding a significant part of the protein backbone to the native conformation. Moreover, we found that there are two distinct knotting mechanisms, which are described here. Also, building on a *loop flipping theory* developed earlier, we present two new theories of protein folding involving the creation and threading of two loops, and explain how our theories can describe the successful folding trajectories for each of the three representative 6_3_-knotted proteins.

## Introduction

Deep learning methods such as AlphaFold (1, 2) provide new tools to study proteins with non-trivial topologies. In particular, AlphaFold has predicted new potential knot types in proteins, including 5_1_, 6_3_ (3, 4), and composite knot types such as 3_1_#3_1_ found in globular and membrane proteins (5). It is possible to spot even more complex knots (e.g. 8_3_) (6), which however should be treated with caution. Such predictions led to the experimental verification of the first ever composite knot, obtained by solving an X-ray structure of the TrmD-Tm1570 protein (PDB ID 8BN1) (7), whose enzymatic activity (the transfer of a methyl group to tRNA) was confirmed (5). In August 2022, a new version, AlphaFold 2 (1, 2), predicted protein structures for 215M sequences that cover almost all known sequences deposited in Uniprot (230M as of 19.10.2022). This data enables researchers to study fundamental questions about protein topology (8, 9), including the complexity of protein topology (9, 10), conservation of protein topology across all species in a given family (11), and the evolution of proteins (12, 13). It also allows investigations into the possible functional roles of knotting in proteins (9, 14, 15), which is mostly unknown except in specific cases (e.g., in the case of TrmD when the knot conformation is necessary to conduct enzymatic activity) (16, 17).

Using experimental methods to determine structure, researchers have identified proteins with knot types 3_1_, 4_1_, 5_2_, 6_1_, and 3_1_#3_1_ (18). Those knots can also occur as slipknots in proteins. A slipknot is created when a knot is formed by part of a protein chain, after which the backbone doubles back so that the entire structure is unknotted (19). The slipknot location, depth of knot (shallow, deep), and the topological fingerprint (which displays all knots that occur in subchains) can be understood using the so-called matrix model (11, 20). The fingerprint analysis shows the existence of both shallow and deep knots in proteins. While shallow knots (i.e., knots where at least one of the tails has fewer than 10 amino acids) can untie through thermal fluctuations, it is more difficult for proteins with much deeper knots to untie (9).

Experimental investigations have shown that some protein knots can self tie in a cell free system and chaperones support folding (21–25). Numerical simulations and theoretical methods further demonstrate that proteins can self tie (10, 26–32), both chaperones (33) and crowding can facilitate knotting (34–36), and ribosomes can play an important role, especially in the case of deeply knotted proteins with attached domains (37, 38).

Such experimental work has led to theories about folding pathways for knotted proteins. In general, numerical simulations suggest that deep knotting is not accidental but takes place in the vicinity of the native position. The knotting pathway arises from the formation of a twisted native loop, followed by either threading the tail, via a slipknot configuration (deeper knot), flipping a twisted loop across the tail, or direct threading. This mechanism (knotting via a slipknot conformation) was also observed in all-atom explicit solvent simulations based on the smallest knotted protein (39, 40). For proteins with a slipknot topology (41), or more complex knots such as 6_1_ (42), the knotting mechanism of flipping a large twisted loop over a smaller loop together with the terminus was suggested, and later a more general *loop flipping* theory of protein knotting was proposed (43) (described in the Results Section).

In this paper, we analyze the 6-crossing knots predicted by AlphaFold 2. In particular, we conducted a comprehensive review of all 215M structures predicted by AlphaFold 2, including over 700K potentially knotted proteins. According to the field of topology, there exist precisely three types of knots with six crossings, denoted by 6_1_, 6_2_, and 6_3_. Through a detailed analysis of the AlphaFold 2 data, we identified new protein families potentially containing a 6_3_ or a 6_1_ knot which are well conserved over a broad range of different species. We then focused on three deeply knotted proteins containing a 6_3_ knot, and found that they can self tie, based on a C*α* coarse-grained structure-based model (44). In addition, we developed an extension of the loop flipping theory that applies to the folding pathways seen in MD simulations of the 6_3_-knotted proteins, and can also be applied to other types of knots. Finally, based on the conservation of knot types in different species we put forward a hypothesis of knot evolution, at least in the case of proteins with six crossings knot types.

### 1. New families of 6-crossing protein knots

Of the 215M structures predicted by AlphaFold 2, we focused on models with pLDDT confidence higher than 70. For each of these, we determined the dominant knot type (45), and found over 700k knotted structures.

Next, we analyzed the proteins with predicted 6 crossings knot types, and classified them based on their potential biological function, sequence similarity, and domain annotations (called architectures by InterPro. Here we will refer to this classification using the InterPro term “families”). Our approach allowed us to check whether the topology is conserved both within each cluster (architecture assignment or similar sequence, when the architecture is unknown) and within the whole family (proteins with the same biological function). As a result, we obtained two additional pieces of information: (i) further verification of a particular knot’s presence in a given organism; and (ii) whether the knot is robustly present (conserved) in a given family (3). Interestingly, these results also singled out the models where, despite good prediction quality, the structures were questionable and led to incorrect topologies.

We then identified *robustly knotted* families, where the knot topology is conserved in the majority of homologous sequences (each family represented by at least 10 members and knotted for more than 50%). Our analysis found a significant number of robustly knotted protein families with 6_3_ and 6_1_ knots (Tab. 2, S1 and Tab. 1, S2 respectively). However, we found only a few single proteins with a 6_2_ knot; and despite using predictions with pLDDT above 70, we identified some problems by visual inspection, see Section B and Tab. S3.

To better understand the quality of the predicted structures, we conducted cross validation using RosseTTaFold and EMSFold (the 600M structures (46)) (different AI approaches (47) for protein structure predictions than AlphaFold) for selected proteins. In general, we found good agreement with RosseTTaFold and poor agreement with EMSFold. In the latter case for proteins predicted as a 6_3_ knot by AlphaFold, the ESMFold predicts a 6_3_ knot in 7% cases, a 5_2_ knot in 20%, and a 3_1_ knot in 65%. There is even bigger difference for the 6_1_ knot type, where X-ray structures are known. For this reason we are not analyzing predictions from EMSFold in detail.

#### A. Proteins with a 6_1_ knot

We found more than 500 proteins with a 6_1_ knot (the dataset of such proteins is in SI). Based on architecture assignment, sequence similarity, fingerprint motif, and careful visual inspection, we identified five robustly knotted families (architectures) with a 6_1_ knot (Tab. 1). Proteins from these families represent over 40 different organisms. The 6_1_ knots are also found in more than 300 proteins without architecture assignment. Using a BLAST search we assigned the majority of these to one of the five families with known architecture. Unclassified proteins with a 6_1_ knot are in a dataset in the SI, and those with fewer than 10 homologous members are in Tab. S2.

The largest family of proteins with a 6_1_ knot is formed based on the *Halocarboxylic acid dehydrogenase DehI* architecture (344 proteins, Tab. 1 and S2), and includes proteins with known X-ray structures (e.g., PDB ID 3BJX) (18). Of these, 7% of homologous proteins predicted by AlphaFold possess a different topology; however based on a visual inspection, this prediction might not be accurate, so classified them as artefact.

All proteins inside this architecture possess the fingerprint motif K6_1_4_1_3_1_ the same as proteins with known X-ray structures (6). However, currently predicted members possess a much deeper knot (especially on the C-terminal) than in the case of known X-ray structures. The loop flipping pathway proposed by (42) (illustrated in the top panel of Fig. 2 in Section 2) should be able to explain the folding of this family, since the longer C-terminal knot tail will not affect knotting in such a case.

The set of proteins with a 6_1_ knot also includes new families whose fingerprint contains a 4_1_ substructure, but where neither a 3_1_ nor a 3_1_ slipknot is observed (Fig. S3). For example, we found that the Papain-like cysteine and Bacteriophage T5 families have a deep 4_1_ subknot. Since these proteins have a shorter C-terminal tail than N-terminal tail, the folding could again follow the loop flipping mechanism (with slipknot threading of the C-terminal) illustrated in Fig. 2. In contrast, in the case of Quinone-dependent D-lactate dehydrogenase where all six members form a 6_1_ knot, both tails possess more than 60 amino acids. In fact, the C-terminal is much longer than the N-terminal. In theory, these proteins could knot by the loop flipping pathway or by the new theories discussed in Section 2, however this will need some numerical verification, since threading 60 amino acids seems unlikely.

**Table 1.** Classification of proteins with at least 10 homologous members forming a 6_1_ knot based on architecture assignment. N of 6_1_ prot. – number of proteins with the 6_1_ knot and the same architecture. % of 6_1_ prot. – percentage of proteins which possess the 6_1_ topology inside the given architecture. Uniprot ID of a representative protein (last column). All proteins with a 6_1_ knot with its knot core location are listed in Tab. S2.

| Domain arch. | N. of<br>6<sub>1</sub> prot. | % of<br>6<sub>1</sub> prot. | Uniprot<br>ID |
| --- | --- | --- | --- |
| 2-haloacid dehalogenase | 180 | 93 | A0A2W0BJ87 |
| Vacuolar protein<br>sorting 62 | 39 | 68 | A0A3M7NTE2 |
| Phage tail collar domain | 25 | 89 | Q3KH70 |
| Papain-like cysteine<br>peptidase | 11 | 73 | D2VLU6 |
| Bacteriophage T5, Orf172<br>DNA-binding | 10 | 91 | A0A3A5VDQ4 |

The other families listed in Tab. 1 and Tab. S2 (which are mainly those with fewer than 10 homologous proteins), are not discussed here; they need further experimental verification.

#### B. Proteins with a 6_2_ knot

We found 41 proteins containing a 6_2_ knot with known and unknown architectures (see Tab. S3). However, we did not find a cluster with more than three members, and those with three members have different topologies. Since there are too few homologous proteins and the 6_2_ knot topology is conserved in less than 20% of homologous structures, it is hard to classify these proteins either as knotted or unknotted. However, it is worth mentioning the case of V4A765-F1, which could form a 6_2_ knot (Fig. S4). The fingerprint includes two 4_1_ knots, and the 6_2_ knot arises from a 4_1_ knot. The knot core of V4A765-F1 covers almost the whole protein backbone, and knot tails are longer than 30 amino acids (thus are probably too long to be packed just by thermal fluctuations). So, this protein will require a more sophisticated pathway, potentially utilizing loop flipping.

#### C. Proteins with a 6_3_ knot

The 6_3_ knot is significant because it is the first prime non-twist knot to occur robustly in a protein family. The first time a 6_3_ knot was detected in a human proteome (in protein called BCSC-1, breast cancer suppressor candidate 1) was based on AlphaFold 1 (3). Herein, taking into account 30 proteomes, we found more than 100 families with at least one member forming a 6_3_ knot. A dataset of these proteins can be found in SI, which includes proteins with known and unknown architectures. Based on architecture assignment, fingerprint pattern motif, and a careful visual inspection, we reduced our focus to 15 robustly knotted families (architectures) with the 6_3_ knot (Tab. 2). We found that these proteins come from over 80 different organisms.

The knot core of these 15 families is formed by two domains, IPR013694 and IPR002035 (based on architecture assignment InterPro/Pfam), connected by linkers (see Fig. 1). The fact that these domains appear together is not a coincidence. IPR013694 – vault protein inter-*α*-trypsin (VIT) and IPR002035 – von Willebrand type A (VWA) are well conserved together and form an inter-*α*-trypsin inhibitor heavy chain (ITIH) (48). Thus all families from Tab. 2 we call ITIH. The knot core of these proteins is surrounded by additional fragments of proteins, either with unknown architecture (the largest group of proteins) or known architecture (domains), which are listed in Tab. 2. The size of these additional domains determines the depth of the 6_3_ knot.

**Fig. 1.**
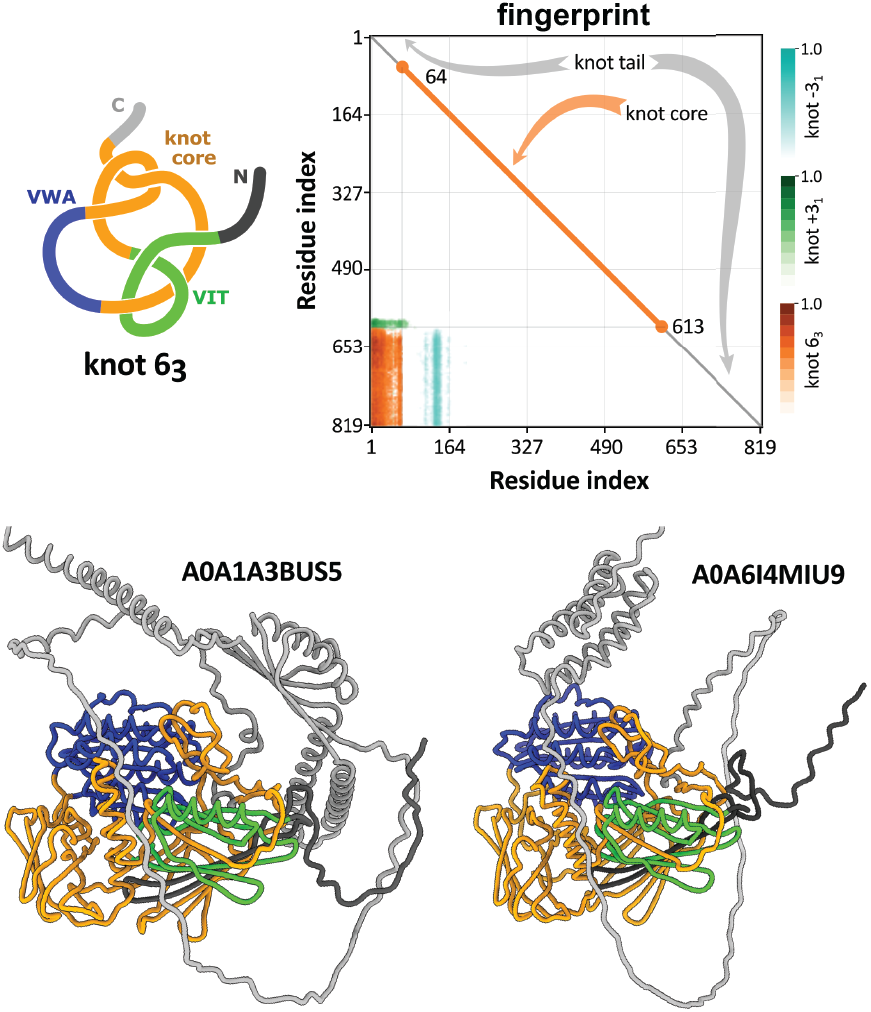
Top left: schematic representation of a protein with 6_3_ knot, from Inter-*α*-trypsin inhibitor heavy chain (called by us ITIH) family, based on A0A6I4MIU9. Draw-ing shows a typical arrangement of the knot core location (orange, green, blue) which include two domains inter-*α*-trypsin (VIT domain, green), and von Willebrand type A (VWA domain, blue). Top right: corresponding fingerprint K6_3_3_1_3_1_ . Bottom: cartoon representation of proteins with a 6_3_ knot used to investigate knotting pathways. The fingerprints and cartoon representation of all representative proteins with a 6_3_ knot are listed in Tab. 2 and shown in SI Fig. S1, S2.

**Fig. 2.**
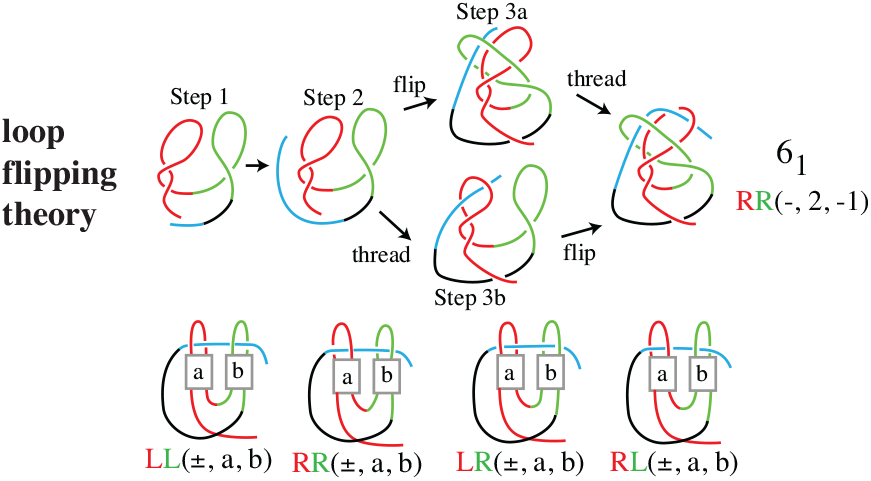
Upper panel: steps of the loop flipping theory. Lower panel: schematic configurations and their notation.

All of the 6_3_-knotted proteins that we investigated are composed of more than 800 amino acids and possess a deep knot, with the length of the tails in the ranges of 49-410 and 46-328, respectively for, the Nand C-terminals. Their fingerprints (shown in Fig. 1 and S1) all have the same general motif in which clipping one terminus produces a 3_1_ knot and clipping the other terminus first produces an unknot and then a +3_1_ knot. On the other hand, even though all of these proteins conserved similar domain architectures, with the same structural cores location, and have the same fingerprint motifs, their sequence similarity ranges from only 20% up to 80%.

Next, using BLAST, we further assigned some proteins with unknown architecture to these 15 families, which are composed of VIT and von Willebrand type A (VWA) domains, which we call ITIH. As a consequence, we found 24 proteins with unknown architecture (Tab. S3), which do not show significant sequence homology among themselves. Since almost all of these proteins form individual clusters and cannot be aligned to other proteins, the conservation of a 6_3_ knot could not be investigated.

Building on the *C*_*α*_ structure-based G*õ* potential (49) and molecular dynamics simulations (44), we found that each protein can self tie into a 6_3_ knot with a success rate of smaller than 2% (Tab. S5). Based on a detailed analysis of these simulations, three pathways can be distinguished (for reaction coordinates we used the number of native contacts (Q) and determined the topology in each frame). Representative kinetics pathways are shown in Figs. 5, 6, 7, as well as in the SI. We explain these pathways in Section 3 after we introduce our theories of knot folding in Section 2.

In order to determine whether a protein can self tie into a 6_3_ knot and how this would occur, we conducted molecular dynamics simulations with structure-based models for three representative proteins which have a low sequence similarity (less, than 25% Tab. S5): O00534 (von Willebrand factor A domain-containing protein 5A, from *human*), A0A1A3BUS5 (Uncharacterized protein, *Mycobacterium asiaticum*, shown in Fig. 1), and A0A6I4MIU9 (VWA domain-containing protein, *Actinomadura physcomitrii*). Due to the large size of the proteins (more than 800 amino acids), we employed a *C*_*α*_ structure-based model which was previously used to study knotted protein folding (41, 42) and was verified in an explicit solvent simulation (39). However, in order to obtain better statistics, we reduced the length of the longer C-terminal tail for A0A6I4MIU9 and A0A1A3BUS5. Details are given in Tab. S5.

**Fig. 3.**
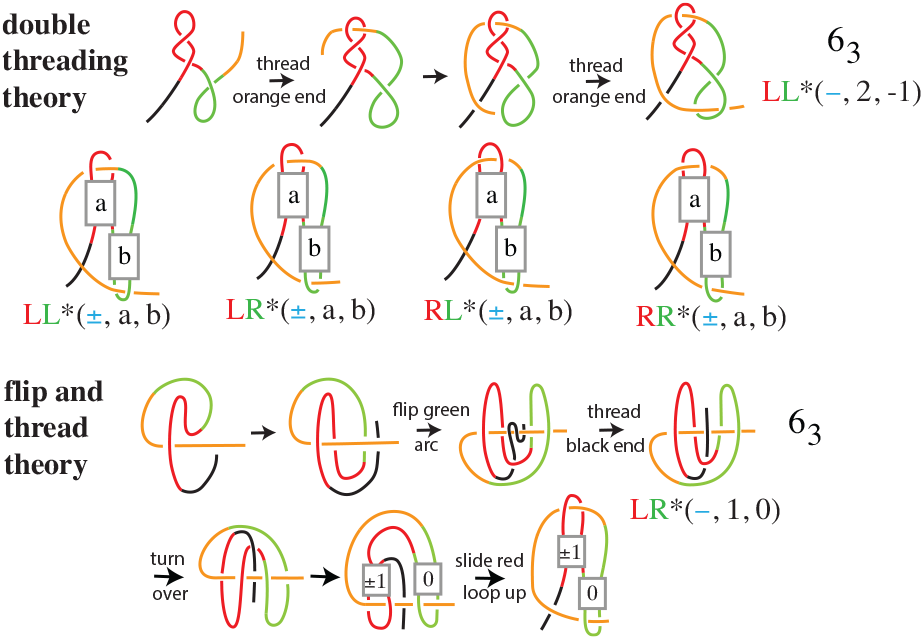
Upper panel: steps of the double threading theory, configurations, and notation. Lower Panel: steps of the flip and thread theory, and demonstration that the configurations produced are the same as those of the double threading theory when *a* = *±*1 and *b* = 0.

**Fig. 4.**
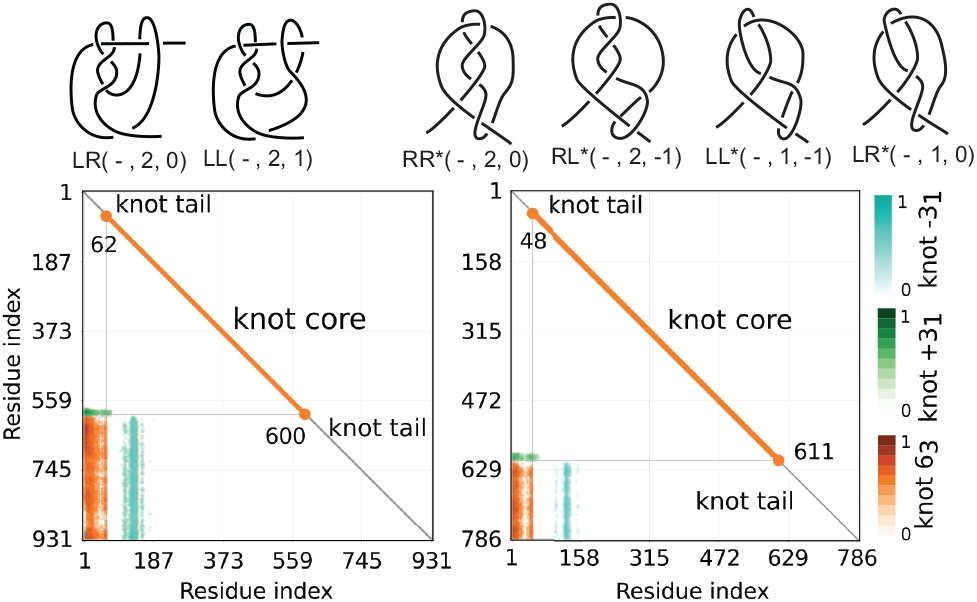
Upper panel: configurations of 6_3_ obtained from our theories whose finger-prints resemble those found with AlphaFold. Lower panel: representative fingerprints for the configurations *LR*(−, 2, 0) and *RR*^*\**^(−, 2, 0).

**Fig. 5.**
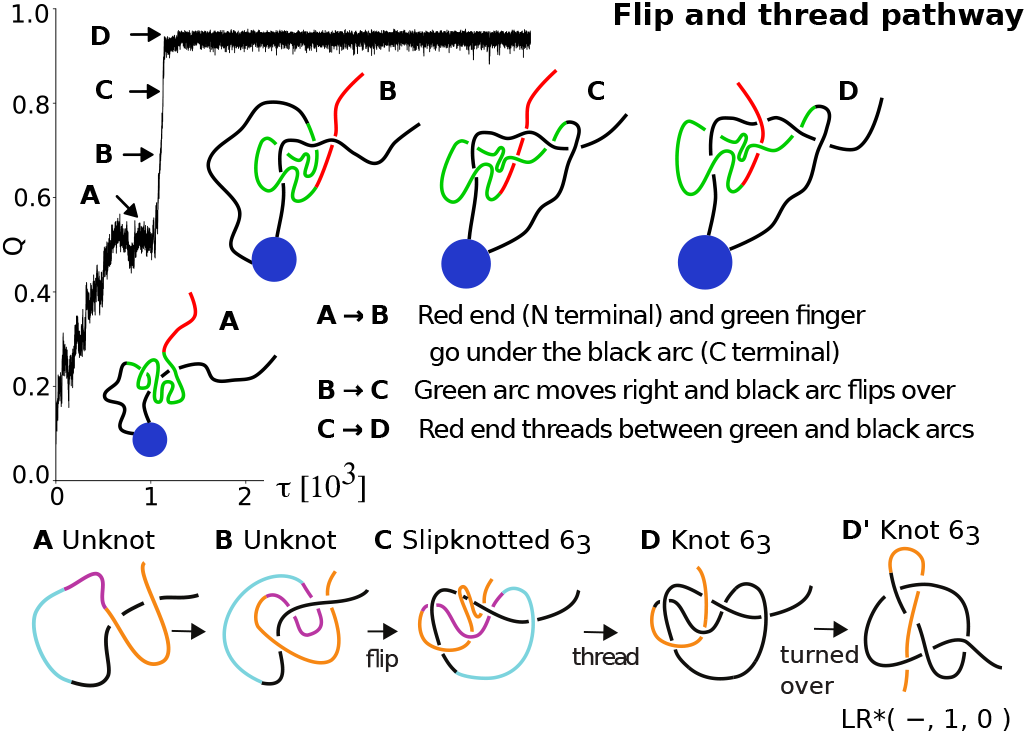
Flip and thread pathway: Example of successful self-tie pathway to a native 6_3_ knot for the O00534 protein. Upper panel: key steps represented via precentage on number of native contacts (Q) versus time (reduce units) based on MD simulations. The green part represents the VIT domain and the blue circle represents the VWA domain (not involved in topology changes). Lower panel: schematic steps showing that O00534 may fold to *LR \** (−, 1, 0) with our flip and thread theory.

**Fig. 6.**
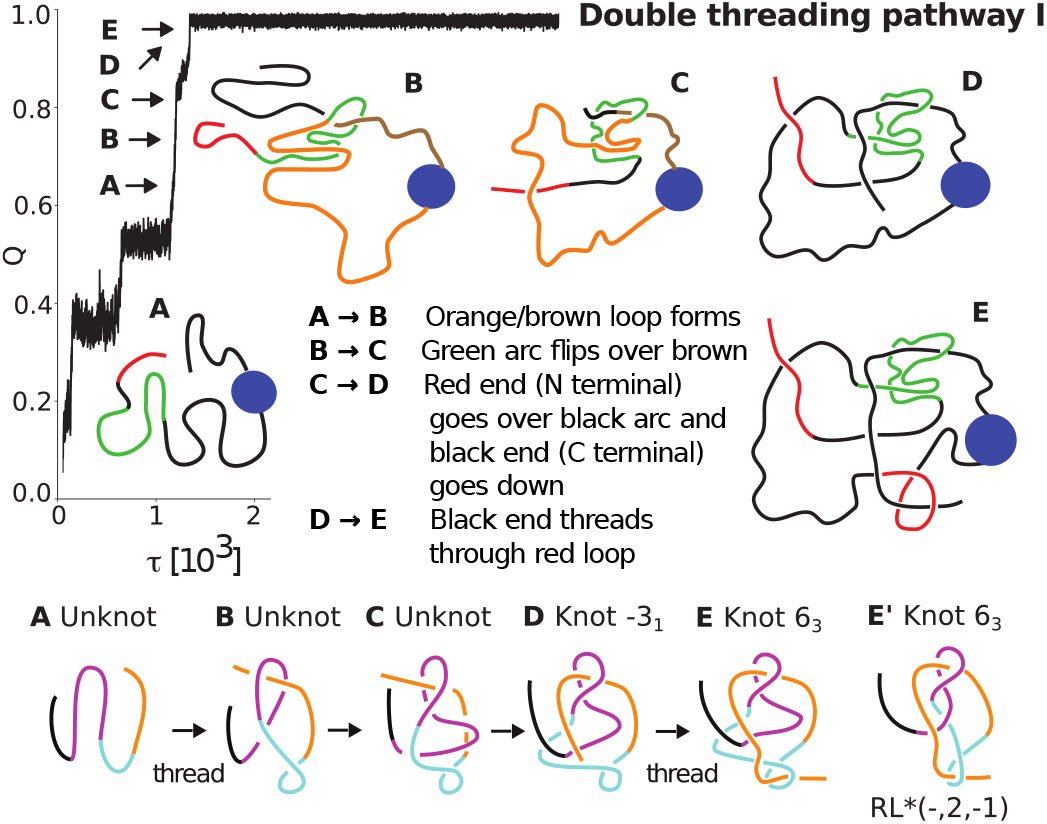
*Double threading pathway I:* Example of a successful self-tie pathway to a native 6_3_ knot for the A0A1A3BUS5 protein. Upper panel: key steps based on MD simulations. The green part represents the VIT domain, and the blue circle represents the VWA domain (not involved in topology changes). Lower panel: schematic steps showing that A0A1A3BUS5 may fold to *RL*^*\**^(−, 2, −1) with our double threading theory.

**Fig. 7.**
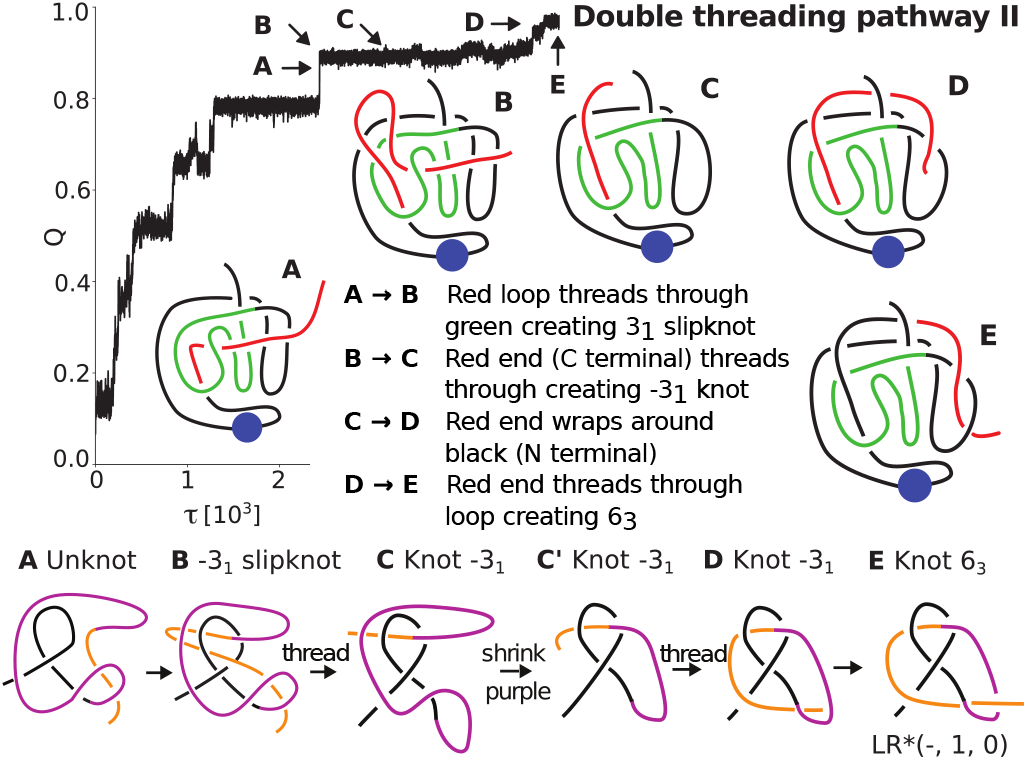
*Double threading pathway II:* Example of a successful self-tie pathway to a native 6_3_ knot for A0A614MIU9. Upper panel: key steps based on a *C*_*α*_ structure-based model with MD simulations. The blue disk represents the VWA domain (not involved in topology changes), and the green part represents the VIT domain. Lower panel: schematic steps showing that A0A614MIU9 may fold to *LL*^*\**^(−, 1, −1) with our double threading theory.

Below we will focus on folding pathways leading to a 6_3_ knot. However, we note that before reaching a 6_3_ knot, the partially-folded protein can get stuck in a 3_1_ knot with a very high number of native contacts established. For A0A6I4MIU9 and A0A1A3BUS5 (with the C-terminal reduced), this occurs in almost 50% of the cases (See Tab. S2). The intermediate state with a 3_1_ knot is structurally very similar to some homological proteins which are predicted by AlphaFold2, Re-seTTaFold, and EMSFold to contain only a 3_1_ knot.

**Table 2.** Classification of protein groups with at least three 6_3_ homologous members based on architecture assignment, Inter-*α*-trypsin inhibitor heavy chain (which we call ITIH) family. Each architecture is composed of VIT and VWA domains connected by linkers (the knot core, Fig. 1), and additional fragments with unknown or known architecture domains (tails). N of 6_3_ prot. – number of proteins with the 6_3_ knot and the same architecture. % of 6_3_ prot. – percentage of proteins which possess 6_3_ topology inside the given architecture. Uniprot ID of a representative protein and position of its knot core (last column). All proteins with 6_3_ knots are listed in Tab. S1.

| Domain arch.<br>attached<br>to VIT;VWA | N. of<br>$6_3$<br>prot. | % of<br>$6_3$<br>prot. | Uniprot<br>ID | knot<br>core |
| --- | --- | --- | --- | --- |
| unclassified<br>domains | 2811 | 52 | A0A6I4MIU9 | 62-618 |
| DUF4349 | 39 | 95 | A0A1A3BUS5 | 59-603 |
| TonB | 19 | 76 | A0A2N3A556 | 86-629 |
| v-SNARE | 16 | 80 | A0A814HQT0 | 49-599 |
| Prot_kinase | 14 | 74 | A0A3D1P8K4 | 410-960 |
| Transl_IF3_C | 6 | 100 | A0A3N6RLM2 | 52-602 |
| FHA | 6 | 100 | B4VXM6 | 181-746 |
| NHL_rpt | 6 | 100 | A0A819SA25 | 53-627 |
| OmpA-like | 5 | 100 | A0A496VSH8 | 86-641 |
| Ankyrin_rpt | 3 | 100 | A0A419ENC1 | 91-636 |
| ART | 3 | 100 | A0A820DFK0 | 206-772 |
| PDZ | 3 | 100 | A0A7V4GDF6 | 311-886 |
| NAD_bd_11;-<br>NAD_bd_2 | 3 | 100 | A0A817RFB0 | 292-864 |
| LPXTG_anchor | 3 | 67 | A0A410VF95 | 114-676 |

### 2. New knot folding theories

Proteins containing a 6_3_ knot are interesting from a theoretical perspective since they cannot fold according to Taylor’s twisted hairpin theory (50), which posits that the chain twists into a loop (the native one) and one terminus threads through the loop to produce a *twist knot*. A knot which results from a twisted hairpin pathway cannot have a knotted intermediate, and clipping one of the threading termini will always produce the unknot. We expect the folding of 6_3_-knotted proteins to follow a more complex theory pathway, because they are not twist knots, have knotted subchains when clipping either terminal Fig. 1, and (in some families) have knotted intermediates. Below, we describe three such potential pathways which involve the threading of two loops, rather than a single loop as in Taylor’s theory.

The first theoretical pathway involving two loops was developed for the 6_1_ knot in DehI by Bolinger, Sulkowska et al. (51), who showed, using molecular dynamics simulations, that *a loop flipping mechanism* could describe its folding pathway. The same mechanism is observed in the case of slipknotted proteins (28). Motivated by the results for DehI, Flapan et al. (43) developed a more general *loop flipping* theory which potentially could describe the folding of all twist knots and a large number of non-twist knots. This theory (illustrated in the top panel in Fig. 2) involves two loops with at most two twists each, where the larger one flips over the smaller one together with a terminal, and the terminal threads through the smaller one. The flipping and threading can occur in either order, as shown in Fig. 2. In the lower panel, we illustrate schematic templates for all possible configurations that can be obtained with this theory and give their notation. See (43) for more details.

We now extend the loop flipping theory to two new theories that produce configurations in which two loops are threaded, but whose steps and final configurations are different from those of the loop flipping theory. The *double threading* theory and the *flip and thread* theory are illustrated in Fig. 3 and described below.

**Steps and notation for double threading theory:**

1. Red and green loops form with *a* and *b* twists respectively, where *|a|, |b|* ≤2 where their sign is the sign of the slope of the overcrossing strand.
2. Orange end threads through red loop, which is designated *L* or *R* according to whether the left or right side of the loop is in front of the orange arc.
3. Orange arc crosses in front or behind black arc, and is designated or +, respectively.
4. Orange end threads through green loop and is designated *L* or *R*, according to whether its left or right side is in front of the arc when rotated 180^?^.

**Steps of the flip and thread theory:**

1. Red-green loop forms.
2. Red-green loop moves down to create a red loop which is threaded by the orange arc, and black end moves up.
3. Black end moves left and green arc flips over/under the rest to create a green loop threaded by orange arc.
4. Black end threads between the red and orange arcs.

On the bottom line of Fig. 3, we turn the final conformation of the flip and thread pathway over, then put the red crossing in a box, and slide the red loop up to see that it produces the same configurations as the double threading pathway when *a* = ±1 and *b* = 0. Note that even though the configurations are the same, the steps of the two theories are different. As a result, the double threading theory can include a knotted intermediate after threading the red loop, while the flip and thread theory never has a knotted intermediate. Nonetheless, in all three theories, threading may occur with the folded ends forming slip knot intermediates, as in the third image of the flip and thread theory.

For the notation, we always list the L or R of the red loop before that of the green loop. Also, the in the notation distinguishes the configurations from the double threading theory and the flip and thread theory from those of the loop flipping theory.

Since we are interested in the folding pathways of 6_3_-knotted proteins, we compared the fingerprints of the configurations from the loop flipping, double threading, and flip and thread theories that produce a 6_3_ knot (listed in Tab. S4) to those of the 6_3_-knotted proteins, and found that only the six shown in the upper panel of Fig. 4 have fingerprints that matched. Representative fingerprints for the configurations *LR*(−, 2, 0) and *RR*^*^(−, 2, 0) are shown in the lower panel, while a typical fingerprint of a 6_3_-knotted protein is shown in Fig. 1. Thus when we analyze potential folding pathways of specific 6_3_-knotted proteins to see if they can be described by one of our theories, we only have to consider the pathways that produce these six configurations.

### 3. Results of structure-based models for folding 6_3_ knot

In order to determine the possible folding pathways and knot folding theories for the representative 6_3_-knotted proteins O00534, A0A6I4MIU9, and A0A1A3BUS5, we did a careful analysis of the successful trajectories based on *C*_*α*_ structure-based model. A more complete visual explanation of the correspondence between the molecular dynamics simulations and our knot folding theories can be found in the SI Fig. S8.

#### Knotting without entangled intermediates, protein O00534

The protein O00534 forms a deep 6_3_ knot, with knot tails of lengths 29 and 176, respectively. Analysis of all knotting pathways (Fig. S5, S6) revealed a folding mechanism to obtain the 6_3_ knot, which is shown in Fig. 5 and described below.

MD simulations showed that folding occurred by packing subdomains, with no intermediate knots and the 6_3_ knot fully formed when almost all native contacts (Q) were established. The key steps are illustrated in the upper panel of Fig. 5 and summarized as follows. Prior to the formation of State A, a loosely-formed conformation that is locally packed in the vicinity of the domains VIT and VWA, shown respectively in green and blue in Fig. 1 and 5, is formed. Then the (red) N-terminal goes under the (black) C-terminal, and the backbone between the N-terminal and the VIT domain forms a (green) finger-like shape which passes under the C-terminal (transition from A to B). Next the green arc slides to the right and the black partially formed VIT domain (with loops connecting the two domains) flips over the VWA domain to form a 6_3_ slipknot (transition from B to C), which corresponds to a Q around 80-90% of contacts. The final step corresponds to threading the N-terminal over the C-terminal to form a 6_3_ knot (transition from C to D). We refer to this folding mechanism as the *flip and thread pathway*.

In the lower panel of Fig. 5, we illustrate the folding steps of the MD simulation schematically, using color to help the reader see which arcs have been moved. To get from State D to D’, we turn the image over so that the orange end is at the bottom and slide the orange loop up. More details are provided in SI Fig. S8. Note that all of the crossings are reversed when we turn the image over. State D’ allows us to identify the configuration as *LR*^*^(−, 1, 0) from our flip and thread theory. There is a close resemblance between the schematic steps in Fig. 5 and the steps of the flip and thread pathway in the lower panel of Fig. 3.

The fingerprint for *LR*^*^(−, 1, 0) also matches that of the protein O00534. Thus our flip and thread theory may describe the folding of the 6_3_ knot in O00534. Further evidence supporting this theory is that we did not see a knotted intermediate in the folding of O00534, which is also the case for the configurations obtained using the flip and thread pathway.

Based on a detailed evaluation of all trajectories (see Tab. S5) we found that it is also possible for the protein O00534 to establish a significant number of native contacts and get stuck in a 3_1_ knot. This corresponds to the intermediates occurring in the two proteins discussed below.

#### Knotting via 3_1_ intermediate, A0A1A3BUS5

The knotting pathway for the protein A0A1A3BUS5 is observed to have a 3_1_ intermediate. The length of the N-terminal of A0A1A3BUS5 is 58 amino acids, whereas the C-terminal is 328 amino acids and is 90% unstructured (moreover, it has a very different conformation in each predicted model based on AlphaFold 2). For our structure-based model, the C-terminal has a very small number of attractive contacts, which are likely random. Thus we reduced the tail to 17 amino acids resulting in a 6_3_ knot on the border between a deep and a shallow knot.

The key steps of the MD simulations for the folding/tying to the native knot of A0A1A3BUS5 protein are illustrated in the upper panel in Fig. 6 and summarized as follows. First, the nucleus of the VWA domain is formed (State A). Next, the black C-terminal creates a loop and starts to interact with the unpacked VIT domain (transition from A to B). Then the VIT domain flips over the VWA domain and a 3_1_ slipknot is formed (transition from B to C) which leads to a 3_1_ knot (transition from C to D). This is the first rate-limiting step (at least in our model). Finally, in the transition from D to E, the C-terminal is folded over and inserted inside the twisted loop to creat a 6_3_ slipknot (the second rate-limiting step) before it pushes through the loop and unfolds to create a 6_3_ knot.

We remark that the red loop in State E was actually formed at State B, but was not included in the drawing for the sake of simplicity. Also, note that the VIT domain is unstructured for a long time and folds only upon interaction with the C-terminal. Thus the middle part of the protein chain is unpacked (no native contacts are observed), and the C-terminal can twist and turn in any space that is unoccupied.

The lower panel gives a schematic illustration of the folding steps, which correspond to the steps illustrated in the top panel of Fig. 3 for the configuration *RL*^*^(−, 2, 1) of our double threading theory. Also, the fingerprint of *RL*^*^(−, 2, 1) corresponds to that of A0A1A3BUS5. Thus our double threading theory may describe the folding of the 6_3_ knot in A0A1A3BUS5.

In summary, during the folding of A0A1A3BUS5, there is an intermediate with a 3_1_ slipknot, followed by a 3_1_ knot, and then the 6_3_ knot is formed when the majority of native contacts are established and the C-terminal has threaded through the twisted loop. We refer to this mechanism as *double threading pathway I*. The analysis of all successful trajectories shows that this is the most typical pathway for a A0A1A3BUS5 protein with reduced size. While this protein can also follow the flip and thread pathway (described above) or double threading pathway II (described below), most often the protein would get stuck in the intermediate state with a native 3_1_ knot and almost 90% of native contacts. The statistics about folding to different types of knots is presented in Tab. S5.

#### Knotting via 3_1_ intermediate, A0A614MIU9

For the protein A0A614MIU9, the N-terminal has 68 amino acids and the C-terminal has 201 amino acids, and it is 90% unstructured. Thus we reduced the C-terminal to 8 amino acids, causing the protein to form a shallow 6_3_ knot. For this reduced protein, we observed the highest number of correctly-folded 6_3_ knots and noted that the folding is well organized with a 3_1_ intermediate (examples of successful knotting are shown in Fig. S7). In 20% of the cases, the protein can establish a high level of native contacts (70%) with the 3_1_ knot in the native position; but in this case, the final threading to obtain a 6_3_ knot is not observed. Statistics are given in Tab. S5.

The key steps of the MD simulations for the folding of the 6_3_ knot in A0A614MIU9 are illustrated in Fig. 7 and summarized as follows. First, the nucleus of the VWA domain forms. Then the nucleus for the VIT domain is established, and the domains start to form the native interface. This leads to establishing 80% of native contacts and the formation of a twisted black loop and a green loop based on the VIT domain (State A). In the transition from A to B, a shallow 3_1_ slipknot is formed when the red loop passes through the green loop. In the transition from B to C, the red C-terminal threads backwards through the green loop, which establishes a 3_1_ knot. This is the first rate-limiting step. Then in a step-by-step manner, the C-terminal begins to form native contacts with an already globular form of the protein (transition from C to D). Finally the C-terminal threads through the black loop and the protein forms a 6_3_ knot (transition from D to E). We refer to this folding mechanism as *double threading pathway II*.

A schematic illustration of the steps of this pathway is shown in the lower panel of Fig. 7 (more details are provided in SI Fig. S8). In this case, we have turned the image in State A over, causing all of the crossings to switch. In the transition from State C to C’ we shrink the purple arc to more clearly reveal that States C’, D, and E correspond to the steps of our double threading theory with configuration *LL*^*^(−, 1, 1). The fingerprint of *LL*^*^(−, 1, 1) also matches that of A0A614MIU9. This suggests that our double threading theory may describe the folding of the 6_3_ knot in A0A614MIU9.

### 4. Discussion

We conducted comprehensive analyses of protein structures predicted by AlphaFold2. We found 700K knotted proteins, with pLDDT score higher than 70. We focused on proteins forming 6-crossing knot types. Based on sequence similarity, architecture organization, and detailed visual inspection, we identified robustly knotted families with the 6_3_ and the 6_1_ knot, where the knot is conserved in the majority of homological structures predicted by AlphaFold 2. We also found a few single proteins forming 6_2_ knots. In the case of 6_1_, and 6_3_ knot types, a knot can be predicted also by the RoseTTaFold approach in the majority of cases.

In the case of proteins with 6_1_ knots, we found families with known topological motif such as 6_1_4_1_3_1_ (including proteins with known X-ray structures, e.g. Halocarboxylic acid dehydrogenase DehI architecture) and new families with K6_1_4_1_ motif, e.g, the Papain-like cysteine and Bacteriophage T5 families.

In the case of proteins with 6_3_ knot types, all predicted structures are composed VIT (vault protein inter-*α*-trypsin) and VWA (von Willebrand type A) domains which form a inter-*α*-trypsin inhibitor heavy chain (ITIH). VIT and VWA domains form the majority of the knot core, which can be surrounded by different biological domains on both terminals, and create a deep 6_3_ knot. We recognized at least 15 different such configurations (80 different organisms), all of which we call ITIH.

Since 6_3_ is the first non-twist knot to be found in protein families and has not previously been analyzed, we focused our analysis of knot folding pathways on proteins containing 6_3_ knots. Proteins with a 6_3_ knot possess more than 800 amino acids, thus we used a *C*_*α*_ coarse-grained structure-based model to explore possible knotting pathways. We found that native contacts are sufficient to guide three proteins (with low sequence similarity) to potential 6_3_ knots in around 2% cases. We found three topologically different pathways, which we explain both from the prospective of native contact formation, and in terms of our double threading and flip and thread theories. In each case, the 6_3_ knot is formed after establishing more than 60% of the native contacts, thus random knotting is not observed in our model.

One could ask how distinct are the pathways we present topologically and from the order of establishing native contacts. If we consider all trajectories which lead to a 6_3_ knot, we find that the folding mechanism generally depends on the depth of the knot, the C-terminal tail of the knot is sufficiently long to form a biological domain, however there are proteins without attached domains which form a shallow knot. In the case of proteins with a deep knot, we observed direct knotting to a 6_3_ knot with no intermediate knot, via the *flip and thread* pathway. Proteins with a shallow knot can also fold via the *flip and thread* pathway, although folding via a 3_1_-knotted intermediate is observed more often, via the *double threading* pathways.

There are at least two pathways for 6_3_-knotted proteins that produce a 3_1_ intermediate, which in our analysis, correspond to *double threading pathways*. The most clear difference is that for *pathway II* (presented based on A0A614MIU9 protein Fig. 7), the VIT domain is able to fold itself by interacting with the chain immediately following this domain. The situation is different for *pathway I* (A0A1A3BUS5 protein Fig. 6), where the VIT domain is unstructured for a long time and folds upon interaction with the C-terminal. This difference in folding allows for much more freedom in the middle of the protein chain, where the terminal can twist and turn in a place that would normally be enclosed. For *pathway II*, two large substructures are formed simultaneously (VIT and VWA domains) and then merge. Thus all topologically significant movements occur only on the surface of the protein.

How is this pathway different from the folding when the 6_3_ knot is established without a 3_1_-knotted intermediate? The *flip and thread* pathway, which leads to a deep 6_3_ knot (as in O00534) looks relatively similar to *the double threading* pathway I (where the VIT domain can be structured only after interacting with the C-terminal). However, in this case the C-terminal tail of the knot can fold locally, forming a stable biological domain. As a result, it is too big to thread into a 3_1_ knot.

The conformation of the protein at this stage can be compared to one which is attached to a surface. One could imagine that when a protein is synthesized on the ribosome, the C-terminal could be just pushed out to make a 3_1_ intermediate and then threaded at the last stage to make a 6_3_. Folding/knotting directly on the ribosome was previously suggested for proteins with a deep knot, e.g., proteins composed of three domains where only the middle one was knotted (37). In this case, external domains folded ahead of the middle one, blocking the threading of either terminal to form a 3_1_ knot.

A protein may not be able to fold into a 6_3_ knot because it gets stuck in a 3_1_ knot. This occurs in around 50% of cases for a shallow 6_3_ knot and around 2% of cases for a deep 6_3_ knot. These 3_1_ knots can be native-like, but there are two options. This could represent a topological trap for the folding of a deeply knotted protein, since a C-terminal could be too long to be threaded into a 6_3_ knot. Here the protein could possibly backtrack (52) to untie. In the case of a structure with a shallow knot, folding can continue and sometimes lead to a 6_3_ knot.

The majority of proteins with a potential 6_3_ knot possess a long C-terminal with an additional domain attached. Thus we hypothesize that in the cell, direct knotting (*flip and thread* pathway, no treading of the C-terminal) or folding on the ribosome could occur. In both cases, knotting could be facilitated by non-native interactions.

However, until the full sequence of any member of the ITIH family has been experimentally solved (all structures deposited in the PDB are too short to form even a 3_1_ knot), the 6_3_ knot should be called a potential knot. Herein, we have shown that it is possible to fold a 6_3_ knot using just native contacts and have established the double threading theory and the flip and thread theory as mathematical models of the pathways involved. These theories extend the loop flipping theory, which we had previously developed for proteins containing non-twist knots.

Note, that some members of the ITIH family are predicted by AlphaFold 2 to contain a deep 3_1_ knot. If so, they probably fold via one of double threading pathways, without final threading of the C-terminal.

Finally, since all members of the ITIH family possess rather deep 3_1_ knots, one could expect that there is a greater likelihood of misfolding in these families, as was observed for proteins with lasso-like type topologies (53). Thus these proteins represent intriguing objects for further investigation.

## 5. Methods

### Dataset

The data was downloaded from the AlphaFold Protein Structure Database (1, 2). It consisted of 215 MM structures. Homologs for each protein were obtained using GGSearch from EMBL-EBI (54) with an e-value cutoff of e-10 and searching within the AlphaFold Protein Structure Database. For the analysis of the conservation of topology in protein families, we clustered the proteins using CD-hit with a 60% sequence identity cutoff (55, 56). For the RoseTTaFold (57) predictions, we used the software available online. In the case of EMSFold (46), the data (around 600M structures) was downloaded (47) and selected protein topologies (after clusturing) were analyzed. We analyzed proteins with pLDDT confidence higher than 70. Evaluation of sequence similarity was performed using the standalone Blast package (58, 59). Clustering was conducted based on CD-HiT. All other bioinformatics analysis was conducted using scripts developed by our team.

### Analysis of all AlphaFold data

The task of detecting knots on such a large scale of over 200M structures is a typical big data challenge. Detecting a knot on a single structure is relatively straightforward thanks to the Topoly Python library (45). However, the scale complicates the analysis, and makes it necessary to develop and use appropriately distributed computing tools to manage the computations and resolve of problems that arise. Among such issues were timeouts due to memory leaks and detection algorithms that occasionally get stuck, problems downloading data due to web server overload or the server becoming unresponsive, etc.

An asynchronous cluster tasks management solution – the kafka-slurm-agent (60), was developed to enable seamless large-scale computations necessary to obtain results for all organisms. Thanks to this solution, the knot detection and identification was computed on three independent Linux clusters managed by slurm. The kafka-slurm-agent has been significantly improved over the first version that was developed during the research work that led to the creation of the AlphaKnot (6) web server and database. The most important extensions included monitoring slurm tasks and termination of timed out jobs as well as enhanced central monitoring of running calculations. The structures were grouped in batches of 4000 each. The knot detection and identification was computed using the Topoly package (45). Information about the proteins was downloaded from the AlphaFold website using dynamic URLs and UniProt, with its API services. To better understand the challenge and scale of the analysis performed we mention some of the parameters: size of all CIF input files: 23 TiB, computing resources used: 3 clusters, 77 nodes, 2004 cores, overall computing time 21 days. In the case of EMSFold, we only analysed proteins which were predicted by AlphaFold 2 to possess a 6 crossing knot.

### The topology detection

All of the models were analyzed by computing the HOMFLY-PT polynomial for 100 random closures; when finding a non-trivial topology, 200 closures were used. The details of the method are explained in (61, 62). The fingerprint method was used to determine the position of the knot core and knot tails based on methods explained in (63). Additionally, knot cores for structures were verified by calculating the Alexander polynomial. A structure is classified as knotted when random closures form a nontrivial knot more frequently than the trivial knot, the pLDDT score for the knot core is above 70, there are no clashes in the knot core region (based on MolProbity tool (64)), the same knot type is found in more than 50% of the protein’s homologs, and no suspicious geometry was found after visual inspection. Structures which did not pass visual inspection and those with obvious problems were rejected from our analysis.

### Molecular dynamics simulations

The proteins were modeled via the C*α* structure-based model with G*õ* potential using MD simulations, as presented in detail in (49). A native contact is considered as formed if the distance between the C*α* atoms is less than 1.2 times their native distance (65). Native amino acid contacts are mimicked by the Gaussian potential (66) based on the SMOG server (44). All simulations were conducted using Gromacs v4.5.4. Leap-frog stochastic dynamics integrator with inverse friction constant equal to 1.0 was used. Time step was equal to 0.0005*τ* . Folding trajectories were executed at T=130 and they started from a number of unfolded and unknotted conformations (200-250, depending on the protein) obtained from long simulations of the proteins at high temperature (T=180). The number of trajectories and types of topologies obtained in the simulations are listed in Tab. S5.

## ACKNOWLEDGMENTS

This work was supported by the National Science Centre #UMO-2018/31/B/NZ1/04016 and 2021/43/I/NZ1/03341 (to JIS), COST EUTOPIA action, the National Science Foundation DMS-1906323 (to HW), and Simons Foundation #822014 (to HW). We also thank Mai Lan Nguyen and Dawid Uchal for their comments and for preparing the drawings.

## References

1. J Jumper, et al., Highly accurate protein structure prediction with alphafold. Nature 596, 583–589 (2021).

2. M Varadi, et al., AlphaFold Protein Structure Database: massively expanding the structural coverage of protein-sequence space with high-accuracy models. Nucleic Acids Res. 50, D439–D444 (2021).

3. AP Perlinska, et al., Alphafold predicts novel human proteins with knots. Protein Sci. p. e4631 (2023).

4. MA Brems, R Runkel, TO Yeates, P Virnau, Alphafold predicts the most complex protein knot and composite protein knots. Protein Sci. 31, e4380 (2022).

5. AP Perlinska, JI Sulkowska, Are there double knots in proteins? prediction and in vitro verification based on trmd-tm1570 fusion from c. nitroreducens. under review 8, e74755 (2022).

6. W Niemyska, et al., AlphaKnot: server to analyze entanglement in structures predicted by AlphaFold methods. Nucleic Acids Res. 50, W44–W50 (2022).

7. F Bruno da Silva, et al., First crystal structure of double knotted protein trmd-tm1570–inside from degradation perspective. bioRxiv pp. 2023–03 (2023).

8. SE Jackson, A Suma, C Micheletti, How to fold intricately: using theory and experiments to unravel the properties of knotted proteins. Curr. opinion structural biology 42, 6–14 (2017).

9. JI Sulkowska, On folding of entangled proteins: knots, lassos, links and ϑ-curves. Curr. Opin. Struct. Biol. 60, 131–141 (2020).

10. WR Taylor, A deeply knotted protein structure and how it might fold. Nature 406, 916–919 (2000).

11. J. Sułkowska, EJ Rawdon, KC Millett, JN Onuchic, A Stasiak, Conservation of complex knotting and slipknotting patterns in proteins. Proc. Natl. Acad. Sci. 109, E1715–E1723 (2012).

12. R Potestio, C Micheletti, H Orland, Knotted vs. unknotted proteins: evidence of knot-promoting loops. PLoS computational biology 6, e1000864 (2010).

13. V Zayats, et al., Slipknotted and unknotted monovalent cation-proton antiporters evolved from a common ancestor. PLoS Comput. Biol. 17, e1009502 (2021).

14. P Dabrowski-Tumanski, A Stasiak, JI Sulkowska, In search of functional advantages of knots in proteins. PloS one 11, e0165986 (2016).

15. SE Strassler, IE Bowles, D Dey, JE Jackman, GL Conn, Tied up in knots: Untangling substrate recognition by the spout methyltransferases. J. Biol. Chem. 298 (2022).

16. T Christian, et al., Methyl transfer by substrate signaling from a knotted protein fold. Nat. structural & molecular biology 23, 941–948 (2016).

17. AP Perlinska, M Kalek, T Christian, YM Hou, JI Sulkowska, Mg2+-dependent methyl transfer by a knotted protein: A molecular dynamics simulation and quantum mechanics study. ACS catalysis 10, 8058–8068 (2020).

18. P Dabrowski-Tumanski, et al., KnotProt 2.0: a database of proteins with knots and other entangled structures. Nucleic Acids Res. 47, D367–D375 (2018).

19. NP King, EO Yeates, TO Yeates, Identification of rare slipknots in proteins and their implications for stability and folding. J. molecular biology 373, 153–166 (2007).

20. KC Millett, EJ Rawdon, A Stasiak, JI Sułkowska, Identifying knots in proteins. Biochem. Soc. Transactions 41, 533–537 (2013).

21. AL Mallam, SE Jackson, Knot formation in newly translated proteins is spontaneous and accelerated by chaperonins. Nat. chemical biology 8, 147–153 (2012).

22. NC Lim, SE Jackson, Mechanistic insights into the folding of knotted proteins in vitro and in vivo. J. molecular biology 427, 248–258 (2015).

23. NP King, AW Jacobitz, MR Sawaya, L Goldschmidt, TO Yeates, Structure and folding of a designed knotted protein. Proc. Natl. Acad. Sci. 107, 20732–20737 (2010).

24. I Wang, SY Chen, STD Hsu, Folding analysis of the most complex stevedore’s protein knot. Sci. reports 6, 1–11 (2016).

25. H Zhang, SE Jackson, Characterization of the folding of a 52-knotted protein using engineered single-tryptophan variants. Biophys. journal 111, 2587–2599 (2016).

26. P Dabrowski-Tumanski, JI Sulkowska, To tie or not to tie? that is the question. Polymers 9 (2017).

27. S Wallin, KB Zeldovich, EI Shakhnovich, The folding mechanics of a knotted protein. J. molecular biology 368, 884–893 (2007).

28. JI Sulkowska, P Sulkowski, J Onuchic, Dodging the crisis of folding proteins with knots. Biophys. J. 96, 81a (2009).

29. JCA Silva, EJF Chaves, GAU de Carvalho, GB Rocha, Investigation of the structural dynamics of a knotted protein and its unknotted analog using molecular dynamics. J. Mol. Model. 28, 1–12 (2022).

30. W Li, T Terakawa, W Wang, S Takada, Energy landscape and multiroute folding of topologically complex proteins adenylate kinase and 2ouf-knot. Proc. Natl. Acad. Sci. 109, 17789–17794 (2012).

31. T Škrbic, C Micheletti, P Faccioli, The role of non-native interactions in the folding of knotted proteins. PLoS computational biology 8, e1002504 (2012).

32. MA Soler, PF Faisca, Effects of knots on protein folding properties. PloS one 8, e74755 (2013).

33. S Niewieczerzal, JI Sulkowska, Knotting and unknotting proteins in the chaperonin cage: Effects of the excluded volume. PloS one 12, e0176744 (2017).

34. J Especial, A Nunes, A Rey, PF Faísca, Hydrophobic confinement modulates thermal stability and assists knotting in the folding of tangled proteins. Phys. Chem. Chem. Phys. 21, 11764–11775 (2019).

35. Y Zhao, P Dabrowski-Tumanski, S Niewieczerzal, JI Sulkowska, The exclusive effects of chaperonin on the behavior of proteins with 52 knot. PLoS computational biology 14, e1005970 (2018).

36. MA Soler, A Rey, PF Faísca, Steric confinement and enhanced local flexibility assist knotting in simple models of protein folding. Phys. Chem. Chem. Phys. 18, 26391–26403 (2016).

37. P Dabrowski-Tumanski, M Piejko, S Niewieczerzal, A Stasiak, JI Sulkowska, Protein knotting by active threading of nascent polypeptide chain exiting from the ribosome exit channel. The J. Phys. Chem. B 122, 11616–11625 (2018).

38. M Chwastyk, M Cieplak, Cotranslational folding of deeply knotted proteins. J. Physics: Condens. Matter 27, 354105 (2015).

39. JK Noel, JN Onuchic, JI Sulkowska, Knotting a protein in explicit solvent. The J. Phys. Chem. Lett. 4, 3570–3573 (2013).

40. S a Beccara, T Škrbic, R Covino, C Micheletti, P Faccioli, Folding pathways of a knotted protein with a realistic atomistic force field. PLoS computational biology 9, e1003002 (2013).

41. JK Noel, J. Sułkowska, JN Onuchic, Slipknotting upon native-like loop formation in a trefoil knot protein. Proc. Natl. Acad. Sci. 107, 15403–15408 (2010).

42. D Bölinger, et al., A stevedore’s protein knot. PLoS Comput. Biol. 6, e1000731 (2010).

43. E Flapan, A He, H Wong, Topological descriptions of protein folding. Proc. Natl. Acad. Sci. 116, 9360–9369 (2019).

44. JK Noel, PC Whitford, KY Sanbonmatsu, JN Onuchic, Smog@ ctbp: simplified deployment of structure-based models in gromacs. Nucleic acids research 38, W657–W661 (2010).

45. P Dabrowski-Tumanski, P Rubach, W Niemyska, BA Gren, JI Sulkowska, Topoly: Python package to analyze topology of polymers. Briefings Bioinforma. 22 (2020) bbaa196.

46. C Hsu, et al., Learning inverse folding from millions of predicted structures in International Conference on Machine Learning. (PMLR), pp. 8946–8970 (2022).

47. RM Rao, et al., Msa transformer in International Conference on Machine Learning. (PMLR), pp. 8844–8856 (2021).

48. M Himmelfarb, et al., ITIH5, a novel member of the inter-alpha-trypsin inhibitor heavy chain family is downregulated in breast cancer. Cancer Lett. 204, 69–77 (2004).

49. C Clementi, H Nymeyer, JN Onuchic, Topological and energetic factors: what determines the structural details of the transition state ensemble and “en-route” intermediates for protein folding? an investigation for small globular proteins. J. molecular biology 298, 937–953 (2000).

50. WR Taylor, Protein knots and fold complexity: some new twists. Comp. Biol. Chem. 31, 151–162 (2007).

51. D Bölinger, et al., A stevedore’s protein knot. PLOS Comput. Biol. 6, 1–6 (2010).

52. S Gosavi, LL Chavez, PA Jennings, JN Onuchic, Topological frustration and the folding of interleukin-1β. J. molecular biology 357, 986–996 (2006).

53. DA Nissley, et al., Universal protein misfolding intermediates can bypass the proteostasis network and remain soluble and less functional. Nat. Commun. 13, 1–16 (2022).

54. F Madeira, et al., Search and sequence analysis tools services from embl-ebi in 2022. Nucleic acids research p. gkac240 (2022).

55. L Fu, B Niu, Z Zhu, S Wu, W Li, CD-HIT: accelerated for clustering the next-generation sequencing data. Bioinformatics 28, 3150–3152 (2012).

56. W Li, A Godzik, Cd-hit: a fast program for clustering and comparing large sets of protein or nucleotide sequences. Bioinformatics 22, 1658–1659 (2006).

57. M Baek, et al., Accurate prediction of protein structures and interactions using a three-track neural network. Science 373, 871–876 (2021).

58. M Silva e Santos, The PhOCoe model–ergonomic pattern mapping in participatory design processes. Work 41 Suppl 1, 2643–2650 (2012).

59. C Camacho, et al., BLAST+: architecture and applications. BMC Bioinforma. 10, 421 (2009).

60. https://github.com/prubach/kafka-slurm-agent. (2022).

61. P Freyd, et al., A new polynomial invariant of knots and links. Bull. Amer. Math. Soc. 12 (1985).

62. JH Przytycki, P Traczyk, Invariants of links of conway type. (2016).

63. KC Millett, EJ Rawdon, A Stasiak, JI Sułkowska, Identifying knots in proteins. Biochem. Soc. Trans. 41, 533–537 (2013).

64. CJ Williams, et al., MolProbity: More and better reference data for improved all-atom structure validation. Protein Sci. 27, 293–315 (2018).

65. PC Whitford, et al., An all-atom structure-based potential for proteins: Bridging minimal models with all-atom empirical forcefields. Proteins: Struct. Funct. Bioinforma. 75, 430–441 (2009).

66. H Lammert, A Schug, JN Onuchic, Robustness and generalization of structure-based models for protein folding and function. Proteins: Struct. Funct. Bioinforma. 77, 881–891 (2009).

