## Supplement for "Proteins containing 6-crossing knot types and their folding pathways"

**Table 1. List of domain architectures of proteins with $6_3$ topology. Each architecture is composed of VIT and VWA domains connected by linkers and potentially with additional domains. For each type, the table provides the number of proteins found with that type, the number of those that are $6_3$-knotted, the percentage of $6_3$-knotted proteins out of all proteins with non-trivial topology with that architecture, and the UNIPROT ID of a representative protein from that family. For the given representative protein the table also gives its knot core range and its length.**

| Domains attached to VIT ; VWA | No. of prot. | No. of $6_3$ prot. | % of $6_3$ prot. | Uniprot ID | Seq. sim. (%) | knot core range | Protein length |
|---|---|---|---|---|---|---|---|
| no extra dom. (only tails) | 5428 | 2811 | 52 | A0A6I4MIU9 | * | 62-618 | 819 |
| DUF4349 | 41 | 39 | 95 | A0A1A3BUS5 | 55 | 59-603 | 913 |
| TonB | 25 | 19 | 76 | A0A2N3A556 | 31 | 86-629 | 777 |
| v-SNARE | 20 | 16 | 80 | A0A814HQT0 | 24 | 49-599 | 834 |
| Prot_kinase | 19 | 14 | 74 | A0A3D1P8K4 | 31 | 410-960 | 1013 |
| Transl_IF3_C | 6 | 6 | 100 | A0A3N6RLM2 | 33 | 52-602 | 883 |
| FHA | 6 | 6 | 100 | B4VXM6 | 34 | 181-746 | 928 |
| NHL_rpt | 6 | 6 | 100 | A0A819SA25 | 26 | 53-627 | 946 |
| OmpA-like | 5 | 5 | 100 | A0A496VSH8 | 32 | 86-641 | 861 |
| Ankyrin_rpt | 3 | 3 | 100 | A0A419ENC1 | 30 | 91-636 | 815 |
| ART | 3 | 3 | 100 | A0A820DFK0 | 22 | 206-772 | 821 |
| PDZ | 3 | 3 | 100 | A0A7V4GDF6 | 33 | 311-886 | 983 |
| NAD_bd_11;– NAD_bd_2 | 3 | 3 | 100 | A0A817RFB0 | 23 | 292-864 | 910 |
| LPXTG_anchor | 3 | 2 | 67 | A0A410VF95 | 33 | 114-676 | 757 |

**Fig. 1.** Visual representation of chosen proteins with $6_3$ topology. For each protein there is noted where VIT and VWA domains are, along with other domains if applicable for protein. Under representation of sequences, in green, there is a visualisation noting where knot core is located, and its starting and ending aminoacids.

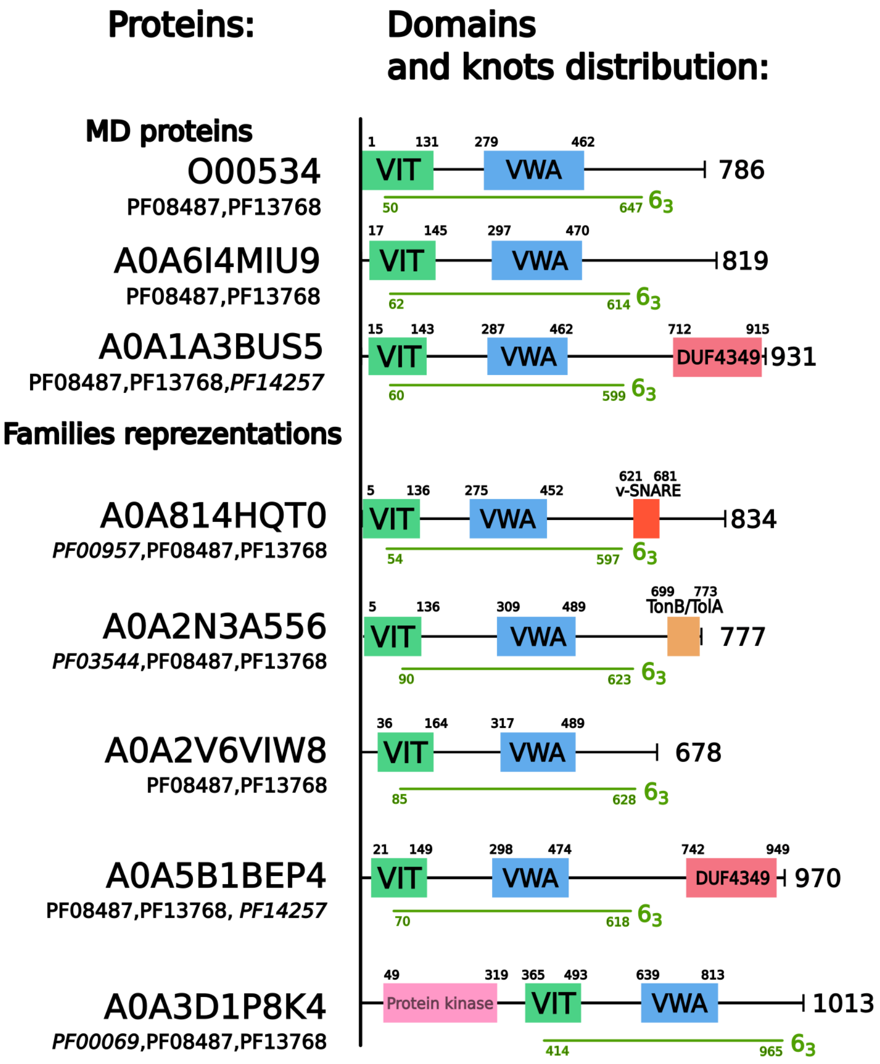

**Fig. 2.** The fingerprint of representative proteins with a $6_3$ knot, based on the Table 1. In this representation of the protein topology, amino acid units are labeled starting at the N-terminus. Each point in the lower left triangle indicates the topology of the subchain starting at the x-axis argument and ending at the y-axis argument. White color denotes the trivial topology $(0_1)$, intensity of the other colors indicates varying probability of a knot type in the given subchain (in accordance with heatmap scales presented on the right of each of the fingerprint matrices). Orange segment on the diagonal shows the range of the knot core within the entire protein, and the remaining grey segments of the diagonal shows the lengths of the knot N- and C-tails.

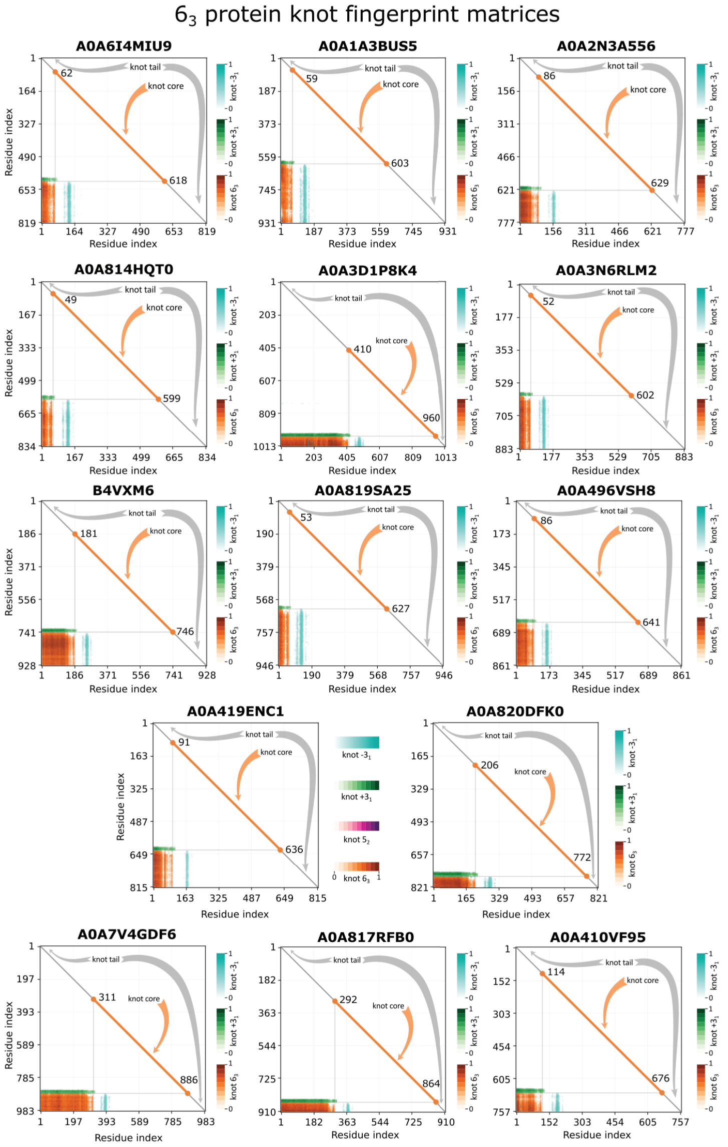

**Fig. 3.** Cartoon representation of example proteins with $6_3$ knot from Table 1. The knot core encompasses the VIT and VWA domains (shown in green and blue, respectively), which are connected via a segment colored in orange. Tails of each knot are shown in grey.

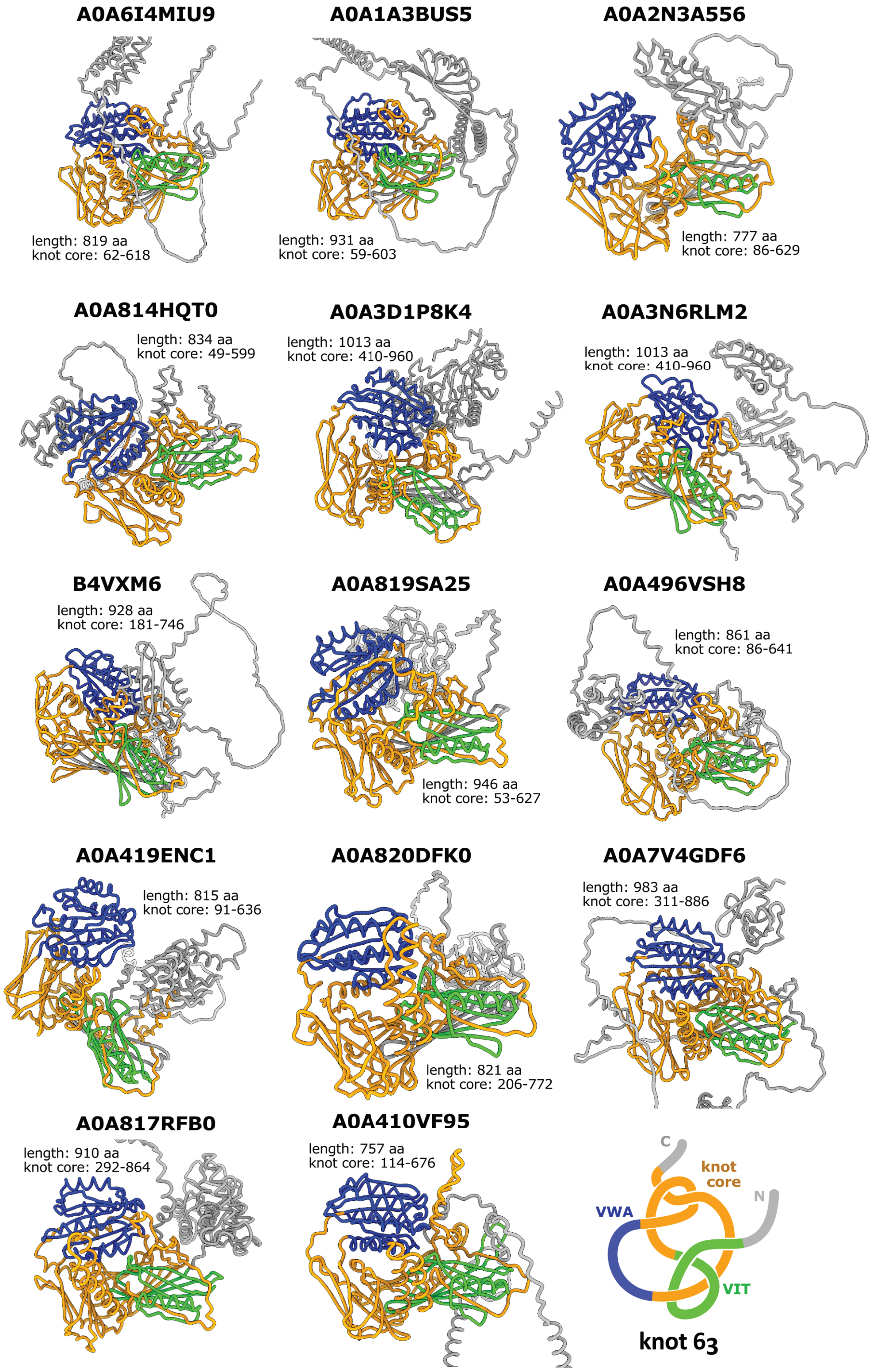

**Table 2. Classification of proteins with $6_1$ topology. For each domain architecture, the table shows the number of proteins that are $6_1$-knotted, the percentage of $6_1$-knotted proteins out of all proteins with non-trivial topology with that architecture, and the UNIPROT ID of a representation protein from that family. For the representative protein, the table also gives its knot core range and its length.**

| Domain arch. | N. of $6_1$ prot. | % of $6_1$ prot. | Uniprot ID | tail knot core tail | Protein length |
|---|---|---|---|---|---|
| 2-haloacid dehalogenase | 180 | 93 | A0A2W0BJ87 | 74-295 | 322 |
| Vacuolar protein sorting 62 | 39 | 68 | A0A3M7NTE2 | 122-516 | 533 |
| Phage tail collar domain | 25 | 89 | Q3KH70 | 328-667 | 817 |
| Papain-like cysteine peptidase | 11 | 73 | D2VLU6 | 288-563 | 569 |
| Bacteriophage T5, Orf172 DNA-binding | 10 | 91 | A0A3A5VDQ4 | 125-333 | 337 |
| Quinone-dependent D-lactate dehydrogenase | 7 | 100 | A0A7S3Q0Z6 | 61-586 | 709 |
| 4Fe-4S dicluster domain-containing protein | 3 | 100 | A0A1W9UJ88 | 18-290 | 692 |
| iron-sulphur binding domain | 3 | 75 | A0A1V5CSN2 | 3-294 | 432 |
| DNA-directed DNA polymerase | 2 | 67 | A0A539EQ13 | 126-876 | 1123 |
| 2OGeDO JBP1/TET, oxygenase domain | 2 | 67 | A0A7K4LXN2 | 276-578 | 655 |
| NB-ARC domain containing protein | 2 | 67 | A0A6L9YXB5 | 699-962 | 1149 |

**Fig. 4.** The fingerprint of representative proteins with $6_1$ knot based on the Table 2.

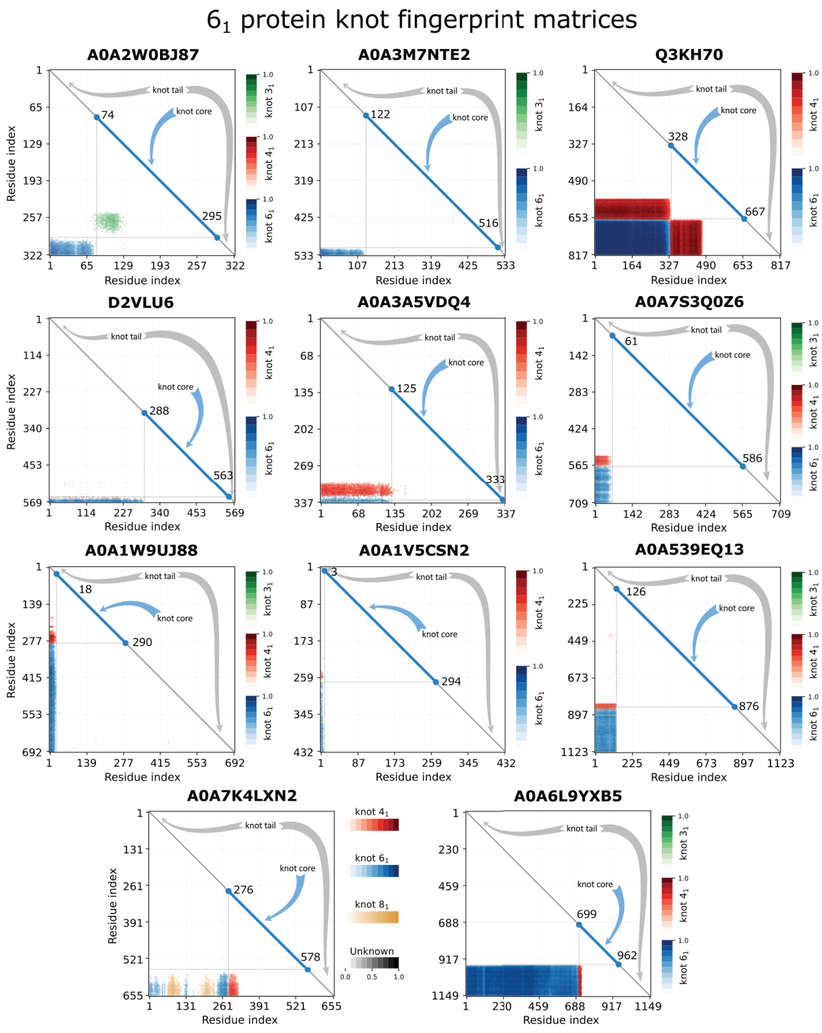

**Fig. 5.** Proteins with potential $6_2$ knot. For proteins with Uniprot IDs V4A765 (top) and A0A845W9E7 (bottom), the figures includes (left) a cartoon representation of the structure with the knot core shown in magenta, and (right) fingerprint of the protein with potential $6_2$ knot, based on the Table 3.

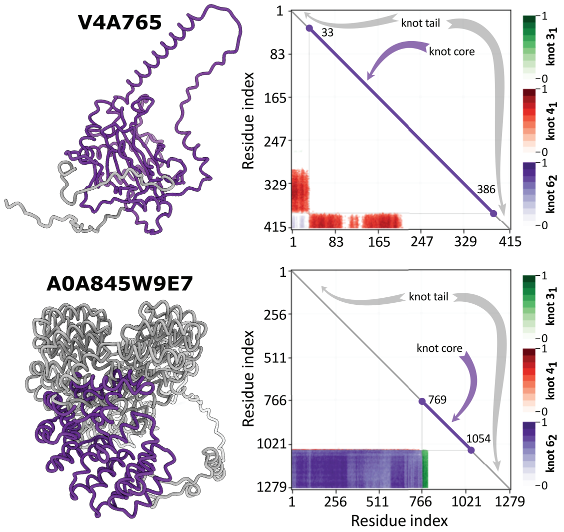

**Table 3. Summary of data with no assigned domain architecture. Architecture was predicted based on a Blast search of sequences of proteins with $6_1$, $6_2$, or $6_3$ knot type, with no architecture as a query and sequences of proteins with known architecture as a subject (architecture information as in Table 1 for $6_3$ and Table 2 for $6_1$). For each protein with no architecture, evaluation was divided into 3 confidence zones based on the lowest E-value found by BlastP as follows— White zone: min. E-value $<$ 0.001; Grey zone: 0.001 $\leq$ min. E-value $<$ 1; Black zone: 1 $\leq$ min. E-value. For each zone the table shows the number of homologous proteins within a zone E-value range which can be assigned to one of the predicted architectures for that knot type, names of the predicted architectures, and the number of proteins which are assigned to that architecture. In brackets next to the predicted architectures, we also note the dominant topology for that architecture (See Table 1 for $6_3$ and Table 2 for $6_1$).**

| Topology | All | White zone | | | Grey zone | | | Black zone | | | Not found by BlastP |
|---|---|---|---|---|---|---|---|---|---|---|---|
| | | N. total | Predicted arch. | N. pred. | N. total | Predicted arch. | N. pred. | N. total | Predicted arch. | N. pred. | |
| $6_1$ | 305 | 195 | DehI ($6_1$)<br>T5_Orf172 ($6_1$)<br>NHL_rpt;ITIH ($6_3$) | 164<br>24<br>1 | 31 | DehI ($6_1$)<br>Collar_fibre ($6_1$)<br>ITIH ($6_3$) | 2<br>1<br>23 | 68 | ITIH ($6_3$) | 3 | 11 |
| $6_2$ | 41 | 1 | NHL_rpt;ITIH ($6_3$) | 1 | 10 | ITIH ($6_3$) | 8 | 25 | | | 5 |
| $6_3$ | 24 | 2 | ITIH ($6_3$) | 2 | 10 | ITIH ($6_3$) | 7 | 10 | | | 2 |
| $3_1\#6_3$, $3_1\#6_1$ | 3 | 0 | | | 0 | | | 0 | | | 3 |

**Table 4. Configurations from the two-loop theories that produce $6_3$ knots**

| **Theoretical pathway** | **Configuration** |
|---|---|
| Loop flipping theory | $LL(-,2,-1)$ |
| (8 configurations) | $LR(-,2,0)$ |
| | $RR(+,1,-2)$ |
| | $RR(+,-2,1)$ |
| | $RL(+,-2,0)$ |
| | $RL(-,0,2)$ |
| | $LR(+,0,-2)$ |
| | $LL(-,-1,2)$ |
| Double threading theory | $RR^{*}(-,2,0)$ |
| (12 configurations) | $RL^{*}(-,2,-1)$ |
| | $LL^{*}(-,1,-1)$ |
| | $LR^{*}(-,1,0)$ |
| | $RL^{*}(-,-1,2)$ |
| | $LL^{*}(-,-2,2)$ |
| | $LR^{*}(+,1,-2)$ |
| | $RR^{*}(+,2,-2)$ |
| | $RR^{*}(+,-1,1)$ |
| | $RL^{*}(+,-1,0)$ |
| | $LR^{*}(+,-2,1)$ |
| | $LL^{*}(+,-2,0)$ |
| Flip and thread theory | $LR_{*}(-,1,0)$ |
| (2 configurations) | $RL_{*}(+,-1,0)$ |

**Table 5. Proteins that were analyzed using Molecular Dynamics simulations. The table shows the Uniprot ID of the three proteins, total number of calculated folding trajectories, number of trajectories which were successful (i.e., simulated protein acquired the $6_3$ knot in the folded state), number of trajectories in which a protein formed a $3_1$ knot, sequence similarity between the O00534 protein and other simulated proteins, and the size of the knot core and N- and C-tails of the knot.**

| Uniprot ID | Total number of trajectories | N. of trajectories $0_1 \rightarrow 6_3$ | N. of trajectories $0_1 \rightarrow 3_1$ | Sequence sim. (%) | Knot core range | Knot N-tail | Knot C-tail |
|---|---|---|---|---|---|---|---|
| O00534 | 1200 | 6 | 8 | * | 30-610 | 29 | 176 |
| A0A6I4MIU9 | 600 | 11 | 228 | 24 | 62-618 | 61 | 8 [a] |
| A0A1A3BUS5 | 500 | 10 | 236 | 26 | 59-603 | 58 | 17 [b] |

[a] For MD A0A6I4MIU9 was cut at its C-terminus to 626 a. a. from its original length of 819 a. a.
[b] For MD A0A1A3BUS5 was cut at its C-terminus to 620 a. a. from its original length of 931 a. a.

**Fig. 6.** Example of folding pathways for protein O00534 presented as Q(t) number of native contacts (left-hand axis) and given topology Topology(t) (right-hand axis).

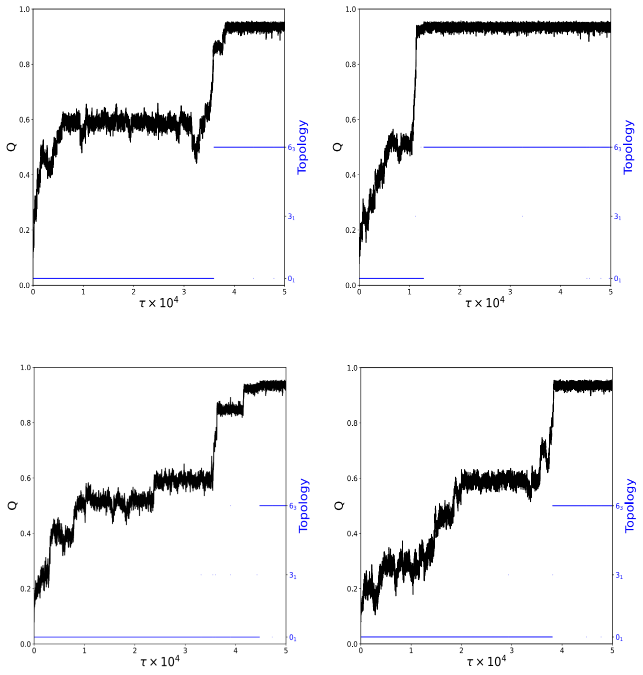

**Fig. 7.** Example of folding pathways for protein A0A6I4MIU9 presented as Q(t) number of native contacts (left-hand axis) and given topology Topology(t) (right-hand axis).

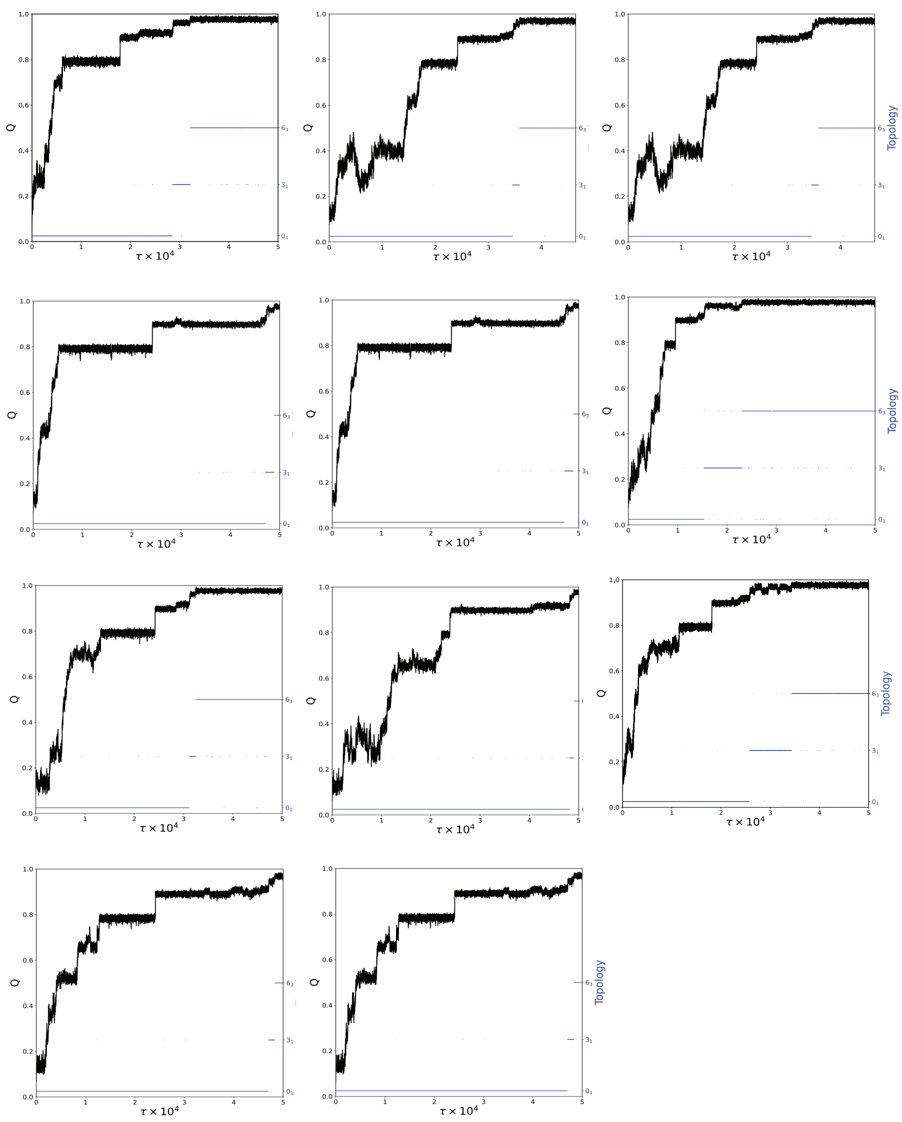

**Fig. 8.** Expanded visual description of representations of protein pathways found on images describing protein folding.

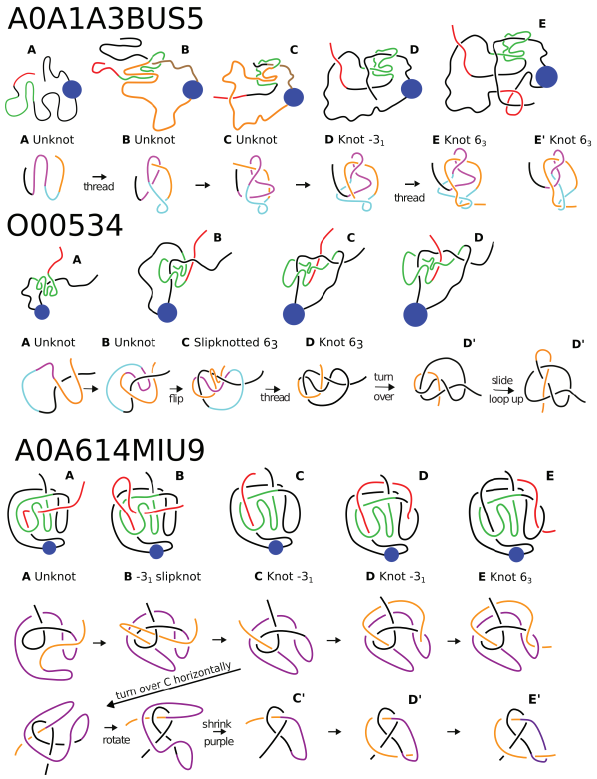

Supplement.xslx/6_3_Raw_Data Full raw data of $6_3$ proteins from 14 domain architectures (Table 1). Provides information about Uniprot, Pfam classification, quality assessment in the form of pLDDT, dominating topology frequency, and additional features such as length and taxonomy.

Supplement.xslx/6_3_Summary Combined statistics for all 14 domain architectures with detailed topology distribution.

Supplement.xslx/6_1_Data Analogous dataset describing proteins with $6_1$ topology from 11 domain architectures. (Table 2).

Supplement.xslx/Classification_conversion Table describing relations between Pfam and InterPro naming notations for both $6_3$ and $6_1$ domain architectures.

Supplement.xslx/ITIH_separated Topological analysis summary of proteins where VIT and VWA domains do not appear together. Additionally, for 23 proteins with VIT domain and the $6_3$ knot, the topological analysis was further extended to include the InterPro domain classification and correct the incomplete and thus potentially confusing results.

Supplement.xslx/Empty_arch Full analysis of proteins with no domain architecture containing knots with 6 crossings. This sheet has 2 parts – raw data and BlastP analysis versus proteins with known architecture (Table 3)

Supplement.xslx/Rosetta Topology comparison between AlphaFold and Rosetta.
